## Supplementary Information for "Learning perturbation-inducible cell states of novel compounds from observability analysis of transcriptome dynamics"

Aqib Hasnain et al.

### 1 Supplementary Text

#### 1.1 Observability maximization for transcriptome dynamics

Here we derive the solution to the observability maximization problem briefly outlined in the Methods section. Recall that we have a state-space representation of the transcriptome dynamics as

$$\begin{aligned} \mathbf{x}_{t+1} &= \mathbf{K}\mathbf{x}_t \\ \mathbf{y} &= \mathbf{W}\mathbf{x}_t \end{aligned} \tag{1}$$

where $\mathbf{x} \in \mathbb{R}^n$ is the (hidden) cell state, $\mathbf{K}$ is the state transition matrix, $\mathbf{W}$ are the unknown gene sampling weights, and $\mathbf{y} \in \mathbb{R}^p$ are the $p$ measurements. The objective, $\mathcal{J}$, is formulated by the signal energy (or output energy) of the system

$$\mathcal{J} = \sum_{i=1}^{m} \mathbf{y}_i^\top \mathbf{y}_i = \sum_{i=0}^{m} \mathbf{x}_0^\top \mathbf{K}^{i^\top} \mathbf{W}^\top \mathbf{W} \mathbf{K}^i \mathbf{x}_0, \tag{2}$$

and we seek the gene sampling weights $\mathbf{W}$ which maximize the objective

$$\begin{aligned} &\max_{\mathbf{W} \in \mathbb{R}^{p \times n}} \mathcal{J} \\ &\text{subject to } \mathbf{W}\mathbf{W}^\top = I_{p \times p}. \end{aligned} \tag{3}$$

The constraint enforces that the rows of $\mathbf{W}$ are orthogonal to each other and that the length of each row be equal to 1. This further avoids the issue of the objective blowing up to infinity. The solution to the above optimization problem is obtained by forming the Lagrangian dual problem and finding the maxima of the the dual objective in terms of the dual variable (a $p \times p$ matrix), $\mathbf{D}$, i.e.

$$\begin{aligned} &\max_{\mathbf{W} \in \mathbb{R}^{p \times n}} \mathcal{J} + \mathcal{L} \\ &\text{where } \mathcal{L} = -\mathrm{tr}\Big((\mathbf{W}\mathbf{W}^\top - I_{p \times p})\mathbf{D}\Big) \end{aligned} \tag{4}$$

and tr() denotes the trace operator. Differentiating the dual objective with respect to $\mathbf{W}^\top$ and equating to 0, we have

$$\begin{aligned} \frac{\partial(\mathcal{J} + \mathcal{L})}{\partial \mathbf{W}^\top} &= \frac{\partial}{\partial \mathbf{W}^\top} \left( \sum_{i=0}^{m} \mathbf{x}_0^\top \mathbf{K}^{i^\top} \mathbf{W}^\top \mathbf{W} \mathbf{K}^i \mathbf{x}_0 - \mathrm{tr}\Big((\mathbf{W}\mathbf{W}^\top - I_{p \times p})\mathbf{D}\Big) \right) \\ &= \frac{\partial}{\partial \mathbf{W}^\top} \left( \sum_{i=0}^{m} \mathrm{tr}(\mathbf{x}_0^\top \mathbf{K}^{i^\top} \mathbf{W}^\top \mathbf{W} \mathbf{K}^i \mathbf{x}_0) - \mathrm{tr}\Big((\mathbf{W}\mathbf{W}^\top - I_{p \times p})\mathbf{D}\Big) \right) \\ &= \frac{\partial}{\partial \mathbf{W}^\top} \left( \sum_{i=0}^{m} \mathrm{tr}(\mathbf{W} \mathbf{K}^i \mathbf{x}_0 \mathbf{x}_0^\top \mathbf{K}^{i^\top} \mathbf{W}^\top) - \mathrm{tr}\Big((\mathbf{W}\mathbf{W}^\top - I_{p \times p})\mathbf{D}\Big) \right) \\ &= \frac{\partial}{\partial \mathbf{W}^\top} \left( \sum_{i=0}^{m} \mathrm{tr}(\mathbf{W} \mathbf{G}^{(i)} \mathbf{W}^\top) - \mathrm{tr}\Big((\mathbf{W}\mathbf{W}^\top - I_{p \times p})\mathbf{D}\Big) \right) \\ &= \frac{\partial}{\partial \mathbf{W}^\top} \left( \mathrm{tr}(\mathbf{W} \sum_{i=0}^{m} \mathbf{G}^{(i)} \mathbf{W}^\top) - \mathrm{tr}\Big((\mathbf{W}\mathbf{W}^\top - I_{p \times p})\mathbf{D}\Big) \right) \\ &= \frac{\partial}{\partial \mathbf{W}^\top} \left( \mathrm{tr}(\mathbf{W}\mathbf{G}\mathbf{W}^\top) - \mathrm{tr}\Big((\mathbf{W}\mathbf{W}^\top - I_{p \times p})\mathbf{D}\Big) \right) \\ &= 2\mathbf{G}\mathbf{W}^\top - 2\mathbf{W}^\top \mathbf{D} = 0 \end{aligned} \tag{5}$$

where the second equality comes from the fact that $\mathcal{J}$ is a sum of $m$ scalars and so applying the trace operator has no effect on the sum, the third equality uses the cyclic property of the trace of products, and the fifth equality uses the fact that $\text{tr}(\mathbf{A}) + \text{tr}(\mathbf{B}) = \text{tr}(\mathbf{A} + \mathbf{B})$. Finally, the Gram matrix, $\mathbf{G}$, is defined to be $\mathbf{G} = \sum_i \mathbf{G}^{(i)} = \sum_i \mathbf{K}^i \mathbf{x}_0 \mathbf{x}_0^\top \mathbf{K}^{i^\top}$, a sum of quadratic forms, which is itself a quadratic form and therefore a symmetric matrix with non-negative, real-valued eigenvalues. From the final equality in Eq. (5) we have

$$\mathbf{G}\mathbf{W}^\top = \mathbf{W}^\top \mathbf{D} \tag{6}$$

which says columns of the eigenvectors of $\mathbf{G}$ are the rows of gene sampling weights $\mathbf{W}$. Moreover, the eigenvector of $\mathbf{G}$ corresponding to the eigenvalue with largest magnitude in $\mathbf{D}$ is the maximizer when $p = 1$.

#### 1.2 Maximizing observability - the integer program

Given the same state-space representation as before, in this section we will develop the mixed-integer program version of the sensor placement framework and highlight the specific relaxations that were made to be able to formulate an analytically tractable optimization problem. We again start by seeking to maximize the signal energy $\mathcal{J}$, however we now add several additional constraints

$$\begin{aligned} &\max_{\mathbf{W} \in \{0,1\}^{p \times n}} \mathcal{J} \\ &\text{subject to } \sum_{i=1}^{n} \mathbf{W}_{ki} = 1,\ k = 1, \ldots, p. \end{aligned} \tag{7}$$

The additional constraints we have added on the optimization problem are i) $\mathbf{W}$ is now a binary matrix, representing whether a gene should be sampled or not, ii) each measurement $y_i$, $i = 1, \ldots, p$ can only be a measurement of a single gene. This problem is clearly an integer programming and in general is difficult to solve for the optimal matrix $\mathbf{W}$. In the section above, we have relaxed both of these constraints, allowing $\mathbf{W}$ to take values in the real numbers and make no restriction on the amount of genes that each measurement can sample from. These relaxations turn the problem from computationally intractable to analytically tractable and the computation time depends largely on an eigendecomposition of $r \times r$ matrices (if using DMD approximations of the network and Gram matrices) or of $n \times n$ matrices with $n$ on the order of 1000 for genetic networks.

#### 1.3 A brief exposition of observability maximization on simulated linear systems

To more clearly emphasize the effect of network topology on the observability maximization problem, in this section we explore the structure of the learned sampling weights as the connectivity of the state-transition matrix, $\mathbf{K}$, is varied for simulated linear systems. We generate fully-connected and sparse block-diagonal matrices to act as the state-transition matrices and for each, we further explore how the number of time points used in the optimization problem (2) affects the learned weights, $\mathbf{w}$.

We simulated 5 distinct network topologies (state-transition matrices) for a simulated 50-dimensional linear system, i) fully-connected, ii) 5 blocks (each fully-connected), iii) 10 blocks, iv) 25 blocks, v) 50 blocks (fully decoupled). For each topology, we obtained the sampling weights that maximize observability for number of time points, $T = 2$ and $T = 5$. Supplementary Figure 1a shows the sampling weights corresponding to each state variable for each topology and time point. For fully-connected topologies, we find that sampling any state variable (gene) for any number of time points leads to similar observability of the entire system, i.e. no one variable contains more information for state reconstruction. As the system transitions from fully-connected to decoupled, key genes are highlighted by higher magnitude weights. The second row in Supplementary Figure 1a corresponds to the learned weights and the state-transition matrices while the third row corresponds to the sorted magnitude of the weights. We can see that as $T$ is increased, the block-diagonal systems' weights transition to more exponential-like curves, resulting in a clear threshold between variables which contribute to observability and variables which do not.

Next, we simulated the same 5 systems as above, however we now set the final (20,20) submatrix to zero. This results in a completely decoupled set of state variables for the last four systems, while for the first system it results in a lack of self edges from the submatrix state variables to themselves. For each topology, we obtained the sampling weights that maximize observability for number of time points, $T = 2$ and $T = 5$. Supplementary Figure 1b shows the sampling weights corresponding to each state variable for each topology and time point. For the first system, we see that lack of self-edges for the lower submatrix results in lower magnitude weights for the corresponding state variables. This discrepancy diminishes as we move from $T = 2$ to $T = 5$ due to information transfer among the genes. For the remaining four systems, we see the same behavior as before, however now the weights corresponding to the state variables with zero submatrices are exactly or nearly zero. As $T$ is increased, we see that the zero submatrix weights converge to zero – information does not transfer to these genes. Initially, for small $T$, the weights are not exactly zero due to nonzero initial conditions. As the initial condition is propagated further in time, its contribution is diminished and the weights converge to zero.

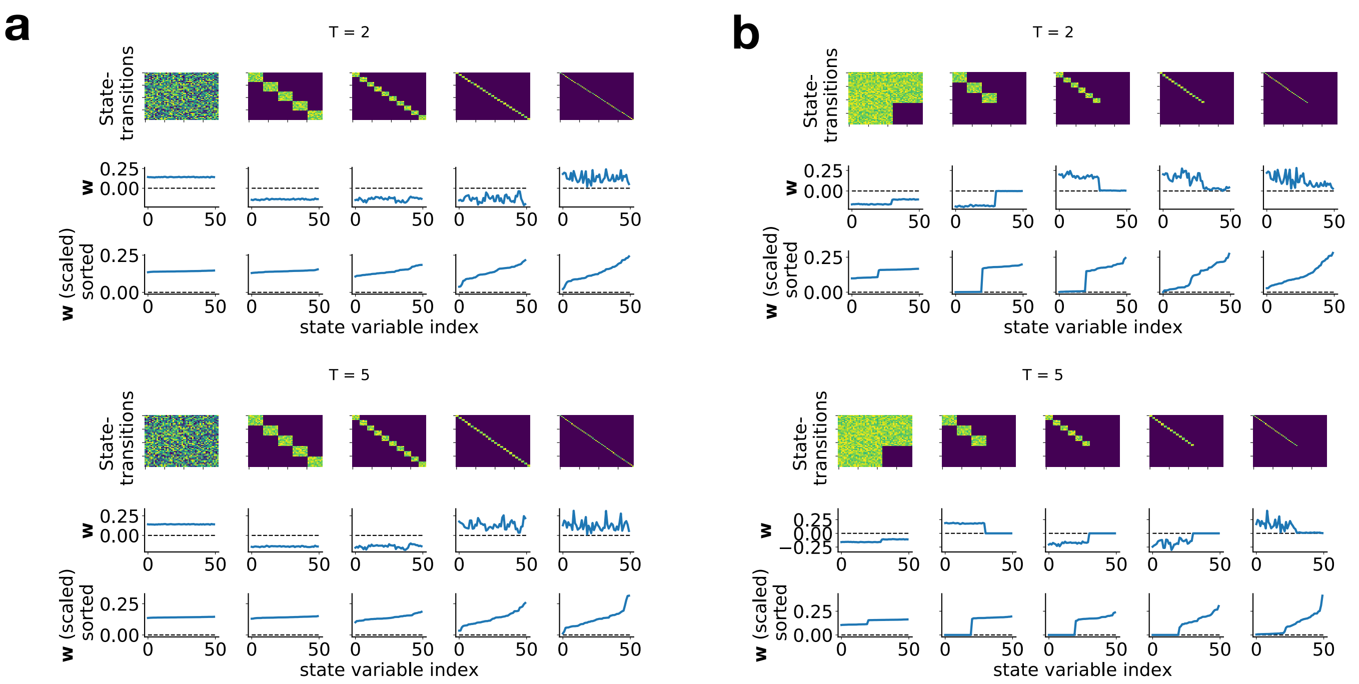


Figure 1: **(a)** From right to left, five systems are simulated and gene sampling weights are obtained and plotted. The system state-transition matrices are shown in the first row, the sampling weights are shown in the second row, and the sorted magnitude of the sampling weights are shown in the third row. **(b)** Same as (a), however now the final (20,20) submatrix of each system is set to zero.

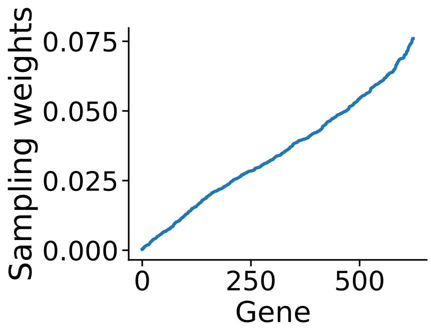


Figure 2: The gene sampling weights that maximize observability for the transcriptomic response of SBW25 to malathion.

For the transcriptomic response of SBW25 to malathion (Supplementary Figure 2), the gene sampling weights are in the regime of nonzero initial conditions for a majority of the genes, nearly fully connected, and relatively large $T$ ($T = 8$). The contribution of each of the above points result in a continuum of sampling weights in which no group of genes is ranked highly compared to the rest of the genes. Notably, even though the learned state-transition matrix is fully connected, we are able to recover a range of sampling weights for the 624 genes. Conversely, for simulated fully-connected networks, we are unable to identify important differences from the sampling weights (the weights are uniformly spread). Consequently, we can conlude that our novel observability analysis is able to capture observable genes from transcriptomic datasets.

#### 1.4 Fold change dynamics of two linear systems

The gene expression dynamics of each experimental condition (negative control and malathion) are well approximated by a linear state-space representation. Specifically, using 10 DMD, we obtained an n-step accuracy of $R^2 = 0.99$ for both conditions. Given this is the case, in this section we want to show when the fold change dynamics of two linear systems can be represented by a linear system. We then define the dynamics as

$$\begin{aligned}\frac{dx_{\text{on}}}{dt} &= ax_{\text{on}} + bu \\ \frac{dx_{\text{off}}}{dt} &= ax_{\text{off}}\end{aligned} \tag{8}$$

where here $x_{\text{on}}$ and $x_{\text{off}}$ are scalar variables for ease of analysis. The variables represent the dynamics in the case where the input is present (*on*) and when the input is absent (*off*), respectively. The input $u$ represents the scalar input of a small molecule, e.g. malathion, that drives the expression of genes in the *on* condition through a step input, i.e. $u(t) = 1$ for all $t > 0$. The solution of the linear ordinary differential equations above are given by

$$\begin{aligned}x_{\text{on}}(t) &= e^{at}x_0 + \int_0^t e^{a(t-\tau)}bu(\tau)d\tau \\ x_{\text{off}}(t) &= e^{at}x_0\end{aligned} \tag{9}$$

where $x(0) = x_0$ for both $x_{\text{on}}$ and $x_{\text{off}}$. We want to show that the fold change response is given by the solution of a linear dynamical system. Taking the fold change of $x_{\text{on}}$ to $x_{\text{off}}$ we have

$$\begin{aligned}x_{\text{fc}}(t) = \frac{x_{\text{on}}}{x_{\text{off}}}(t) &= 1 + \int_0^t e^{-a\tau}\frac{b}{x_0}d\tau \\ &= 1 + \frac{b}{ax_0} - \frac{b}{ax_0}e^{at} \\ &= 1 + \alpha - \alpha e^{at}.\end{aligned} \tag{10}$$

To show that there exists a linear ordinary differential equation (ODE) that gives rise to the above solution $x_{\text{fc}}(t)$, we apply the steps to solve linear ODEs using integrating factors but in reverse order. We know in advance that the integrating factor should take the form $e^{at}$ and we start by dividing both sides of (10) by this integrating factor

$$e^{-at}x_{\text{fc}} = e^{-at}(1+\alpha) - \alpha. \tag{11}$$

We next differentiate both sides and integrate both sides with respect to $t$

$$\int_0^t \frac{d}{dt}\Big(e^{-at}x_{\text{fc}}\Big)dt = \int_0^t ae^{-at}dt - \int_0^t \alpha ae^{-at}dt, \tag{12}$$

then once again differentiating both sides gives

$$\frac{d}{dt}\Big(e^{-at}x_{\text{fc}}\Big) = ae^{-at} - \alpha ae^{-at}. \tag{13}$$

Applying the product rule to the left hand side, we have

$$\begin{aligned}e^{-at}\frac{dx_{\text{fc}}}{dt} - ae^{-at}x_{\text{fc}} &= ae^{-at} - \alpha ae^{-at} \\ &= e^{-at}(a - \alpha a).\end{aligned} \tag{14}$$

Finally, multiplying through by the integrating factor, $e^{at}$, and solving for $\frac{dx_{\text{fc}}}{dt}$, we obtain

$$\frac{dx_{\text{fc}}}{dt} = ax_{\text{fc}} + a - \alpha a \tag{15}$$

which is a linear first order ODE, i.e. a linear dynamical system with a step input and $\alpha = \frac{b}{ax_0}$. The importance of this result is to to be able to say that if the dynamics of the transcriptome in each experimental condition are well represented by a linear system, than the fold change dynamics, under the stated assumptions, can also be well represented by a linear system.

We briefly remark on the extension to the multivariate case. Under the assumption that the system dynamics, $A$, is diagonalizable, the above analysis holds. One such transformation which diagonalizes the the dynamics is given by the set of eigenvectors of $A$. Formally, if we now have system dynamics with state, $\mathbf{x} \in \mathbb{R}^n$, such that

$$\begin{aligned}\frac{d\mathbf{x}_{\text{on}}}{dt} &= A\mathbf{x}_{\text{on}} + Bu \\ \frac{d\mathbf{x}_{\text{off}}}{dt} &= A\mathbf{x}_{\text{off}},\end{aligned} \tag{16}$$

applying the transformation $\tilde{\mathbf{x}} = T^{-1}\mathbf{x}$, where $T \in \mathbb{R}^{n \times n}$ is the matrix of eigenvectors of $A$, results in the transformed systems

$$\begin{aligned} \frac{d\tilde{\mathbf{x}}_{\text{on}}}{dt} &= D\tilde{\mathbf{x}}_{\text{on}} + \tilde{B}u \\ \frac{d\tilde{\mathbf{x}}_{\text{off}}}{dt} &= D\tilde{\mathbf{x}}_{\text{off}}, \end{aligned} \tag{17}$$

where $\tilde{B} = T^{-1}B$. To solve for the fold change dynamics in the multivariate case, we cast the state coordinates into a diagonal matrix, i.e. $\text{diag}(\tilde{\mathbf{x}})$, and compute $\text{diag}(\tilde{\mathbf{x}}_{\text{on}})(\text{diag}(\tilde{\mathbf{x}}_{\text{off}}))^{-1}$. Since the solution in each coordinate is uncoupled from other coordinates, we then have $n$ solutions, each as in Eq. (10).

The case where the above derivation does not hold when the eigenvalues of $A$ have zero real part, i.e. they are exactly zero or have purely sinusoidal response (corresponding to periodic orbits). In this case, the fold change in the coordinate corresponding to zero eigenvalues will approach infinity or it will not be possible to represent the fold change dynamics as a sum of weighted exponentials, e.g. $\tan(x)$. However, such a case would be improbable in a data-driven application for gene regulatory networks. Moreover, any eigenvalue with magnitude zero corresponds to fast dynamics that decay quickly, not significantly contributing to the overall dynamics. There is a time-scale separation between low magnitude and high magnitude eigenvalues, and low magnitude eigenvalues may be removed with little to no effect on the dynamics.

### 2 Supplementary Figures

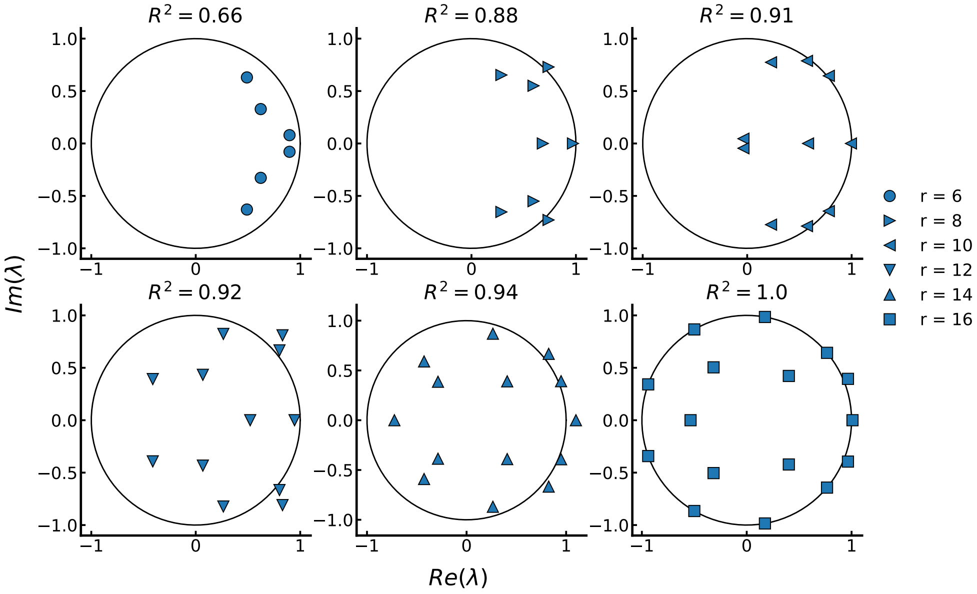


Figure 3: The eigenvalues of the DMD operator plotted in the complex plane for varying number of modes.

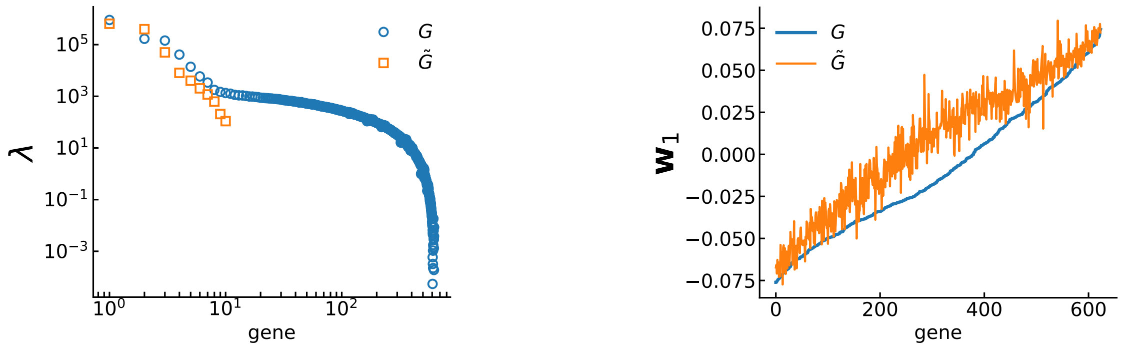


Figure 4: (Left) Approximation of the eigenvalues of the Gram matrix by the reduced order model given by DMD. The full Gram matrix eigenvalues are given in blue circles and the reduced Gram matrix eigenvalues are given in orange squares. (Right) Approximation of the leading eigenvector of the Gram matrix by the reduced order model given by DMD. This eigenvector corresponds to the gene sampling weights in the main text.

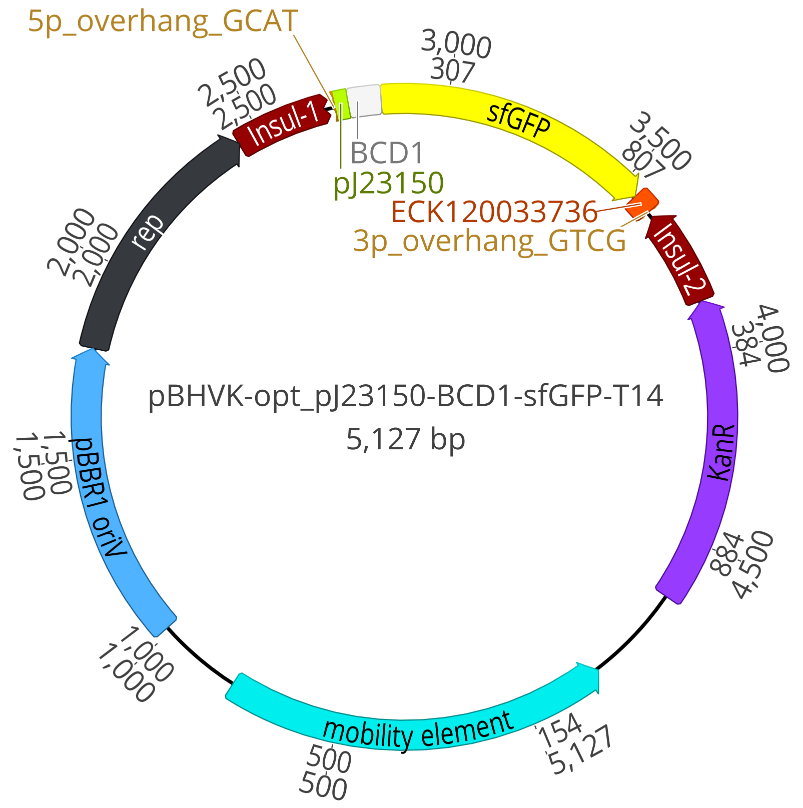


Figure 5: The full plasmid map of pBHVK with the reporter cassette. The two BsaI cut sites on either side of the promoter (5p and 3p overhangs), pJ23150, are used in Golden Gate Assembly to replace the promoter sequence with a promoter used for malathion sensing. A bicistronic design is used for the ribosome binding site, BCD1. A terminator from the set of Voigt lab terminators is used, ECK120033736. For fluorescent reporting, super folder GFP (sfGFP) is used. See Table 3 for sequences of the overhangs, terminator, ribosome binding site, and sfGFP. See Table 2 for sequences of the promoters used in the reporter library.

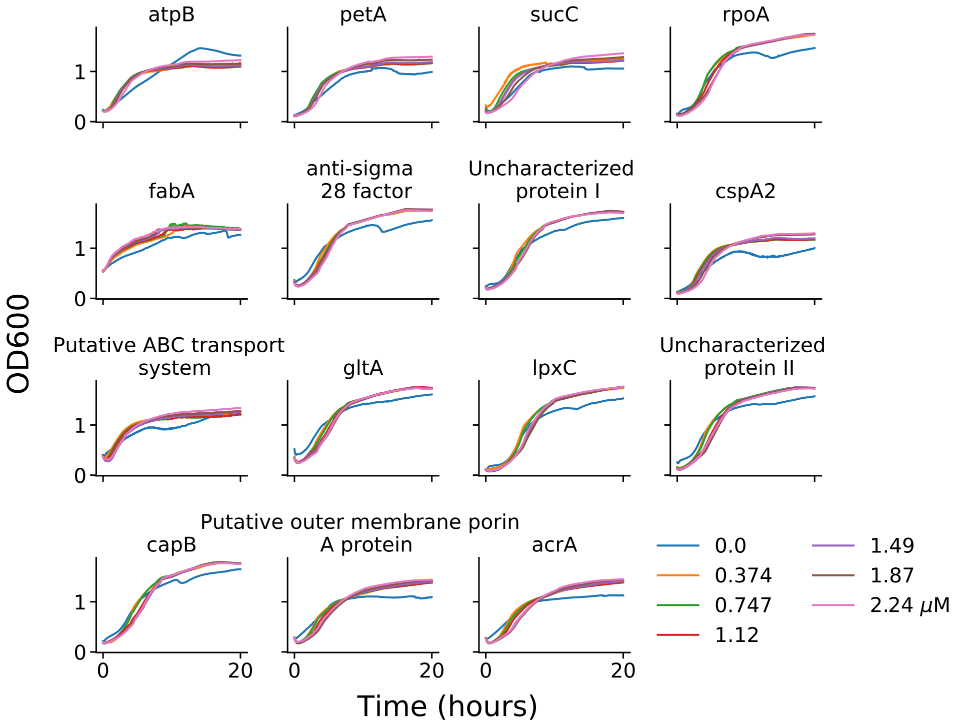


Figure 6: Growth curves of each malathion reporter subject to malathion induction by means of Spectracide.

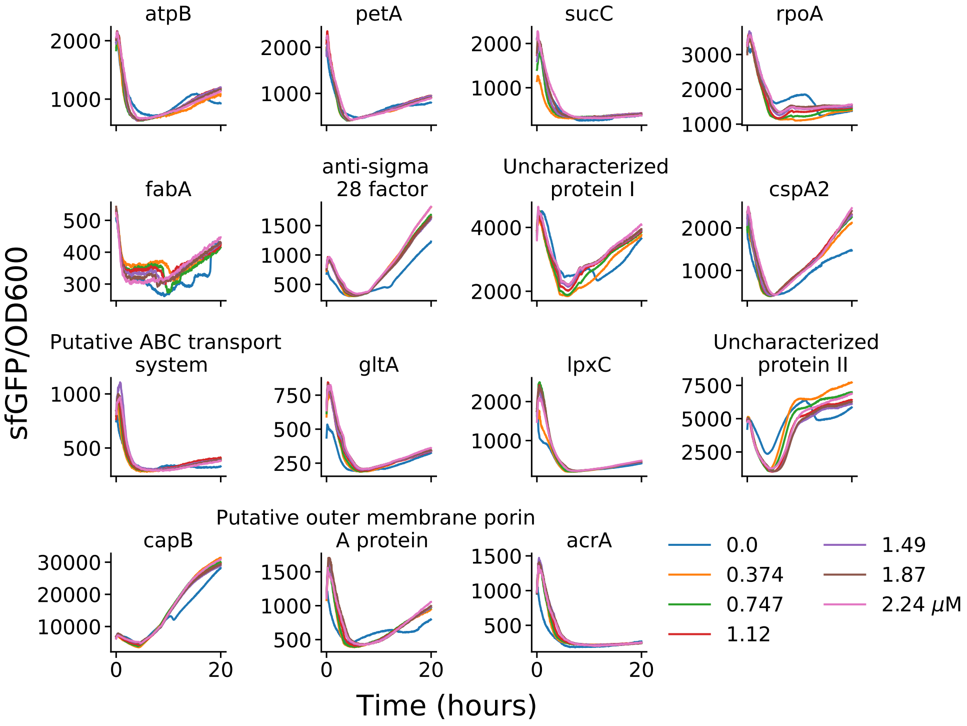


Figure 7: OD normalized sfGFP fluorescence (arbitrary units) of each malathion reporter subject to malathion induction by means of Spectracide.

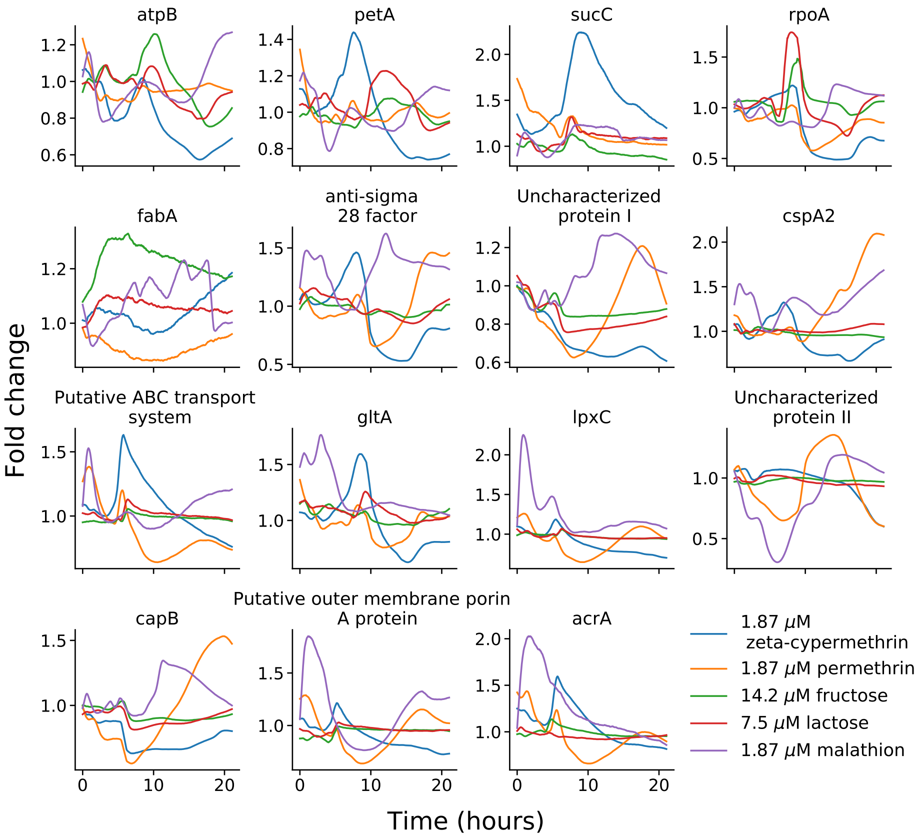


Figure 8: The fold change response of the reporters subjected to five distinct compounds. The fold change is taken as $(\text{sfGFP/OD})_{\text{compound}}/(\text{sfGFP/OD})_{\text{control}}$.

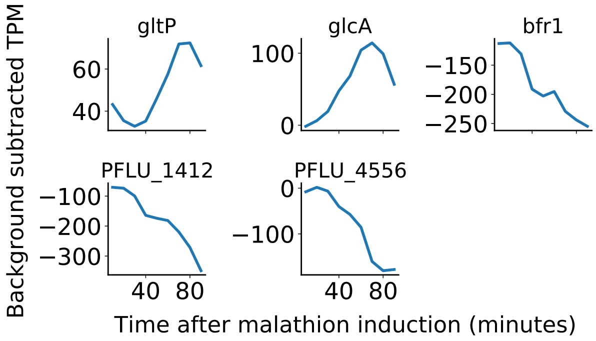


Figure 9: The five differentially expressed genes called by DESeq2 after multiple-testing correction with the Benjamini-Hochberg procedure.

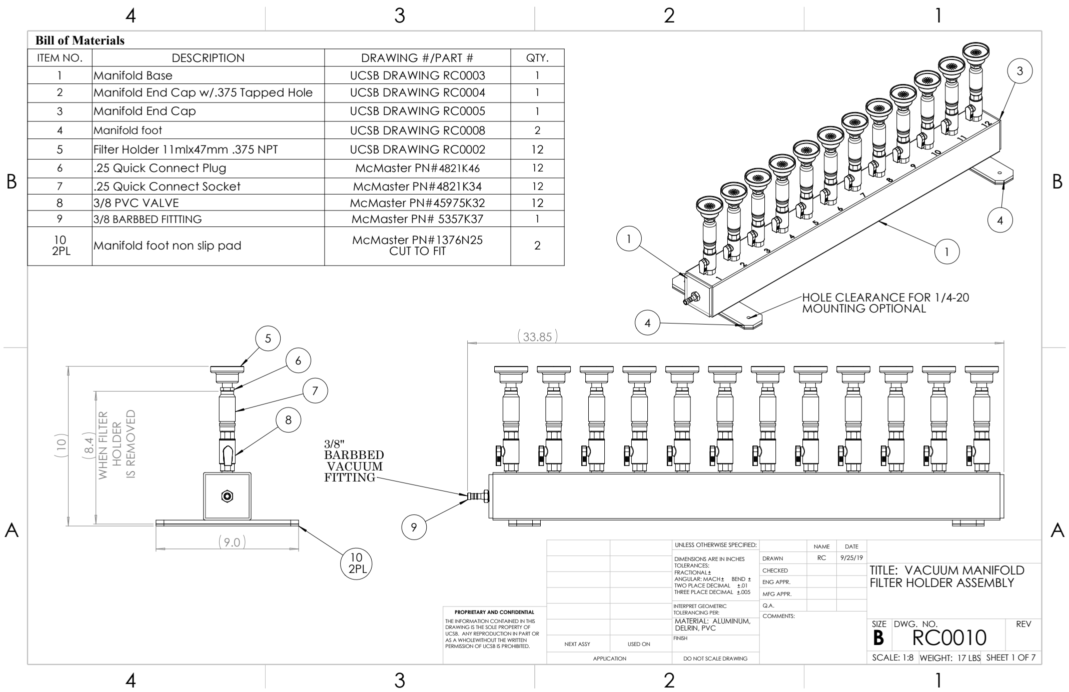


Figure 10: Vacuum manifold design for rapid sampling of mRNA dynamics.

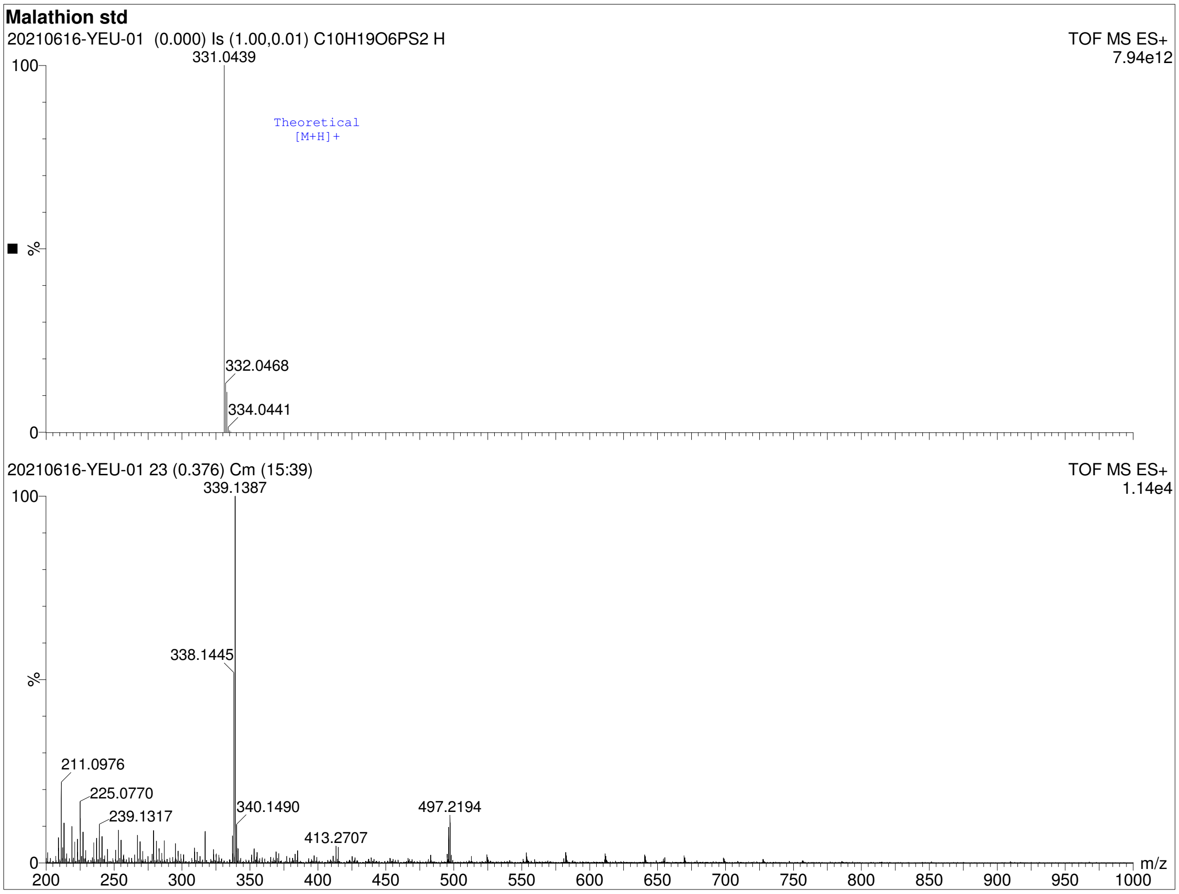


Figure 11: Mass spectrum of malathion (Millipore Sigma Catalog no. 36143) given by time-of-flight mass spectrometry. The theoretical mass spectrum is shown in the upper spectrum and the measured mass spectrum is shown in the lower spectrum.

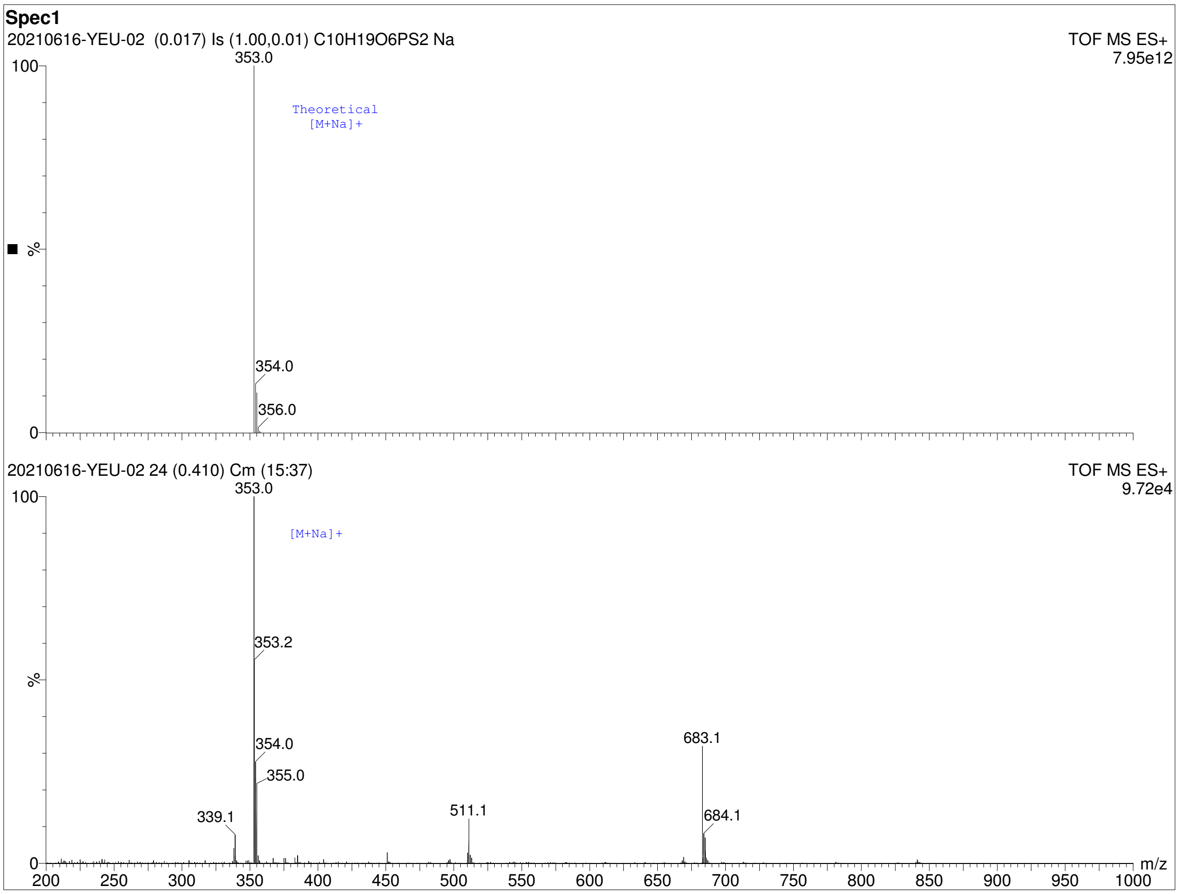


Figure 12: Mass spectrum of Spectracide (replicate 1) (Spectracide Catalog no. 071121309006) given by time-of-flight mass spectrometry. The theoretical mass spectrum of malathion is shown in the upper spectrum and the measured mass spectrum of Spectracide is shown in the lower spectrum.

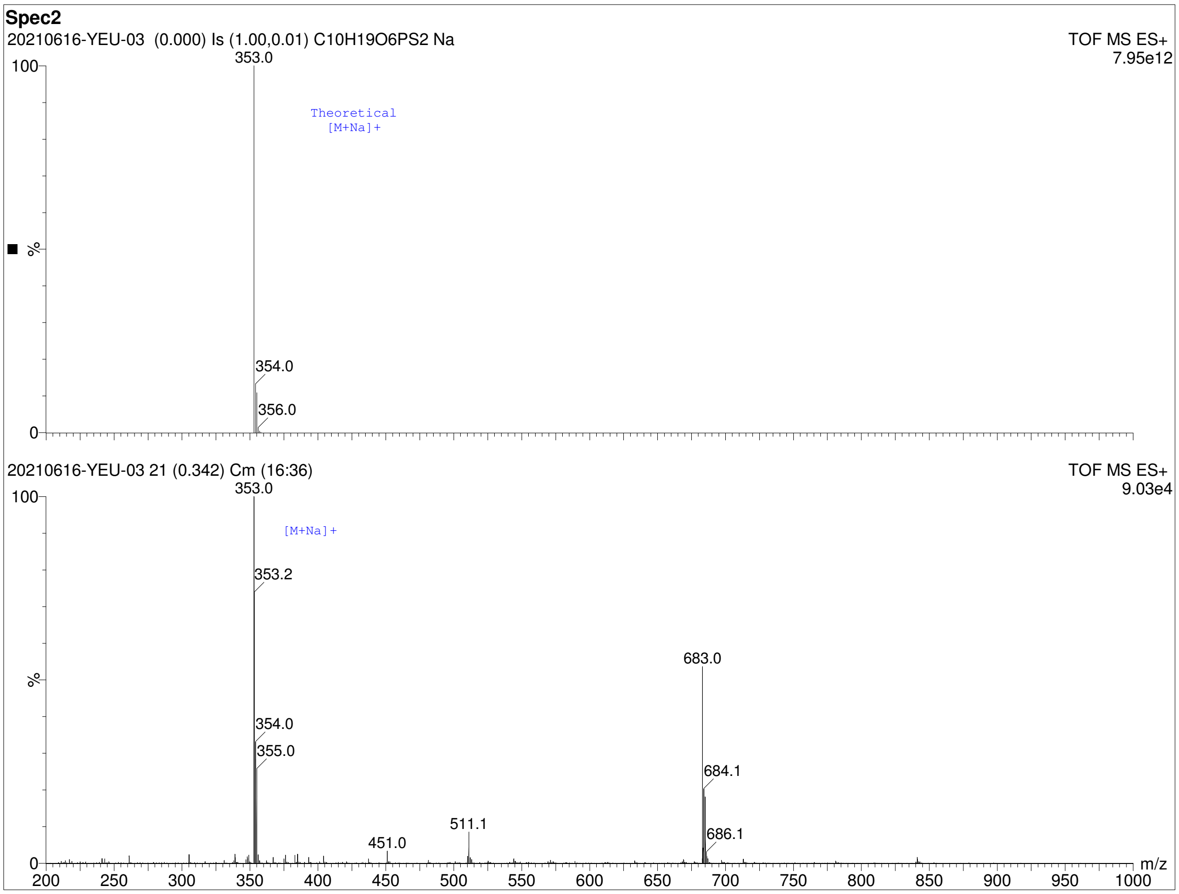


Figure 13: Mass spectrum of Spectracide (replicate 2) (Spectracide Catalog no. 071121309006) given by time-of-flight mass spectrometry. The theoretical mass spectrum of malathion is shown in the upper spectrum and the measured mass spectrum of Spectracide is shown in the lower spectrum.

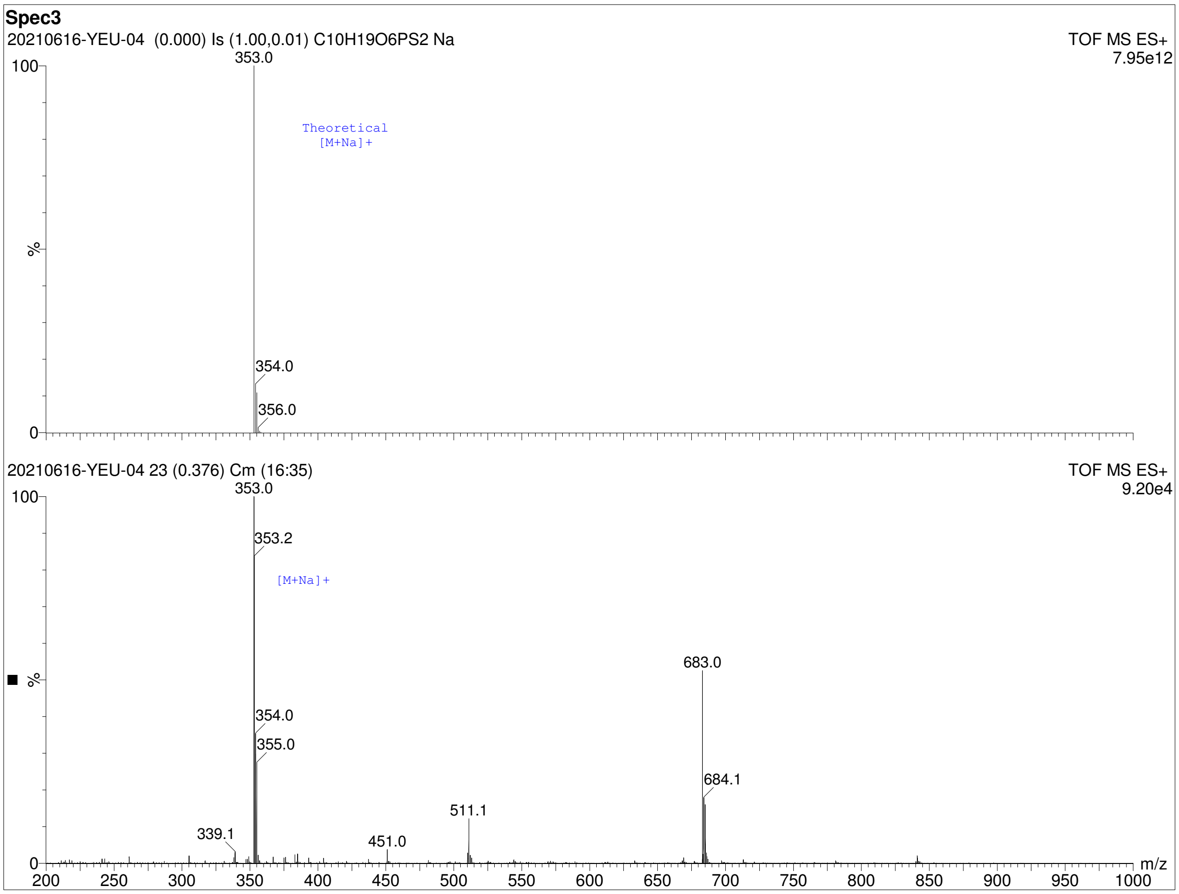


Figure 14: Mass spectrum of Spectracide (replicate 3) (Spectracide Catalog no. 071121309006) given by time-of-flight mass spectrometry. The theoretical mass spectrum of malathion is shown in the upper spectrum and the measured mass spectrum of Spectracide is shown in the lower spectrum.

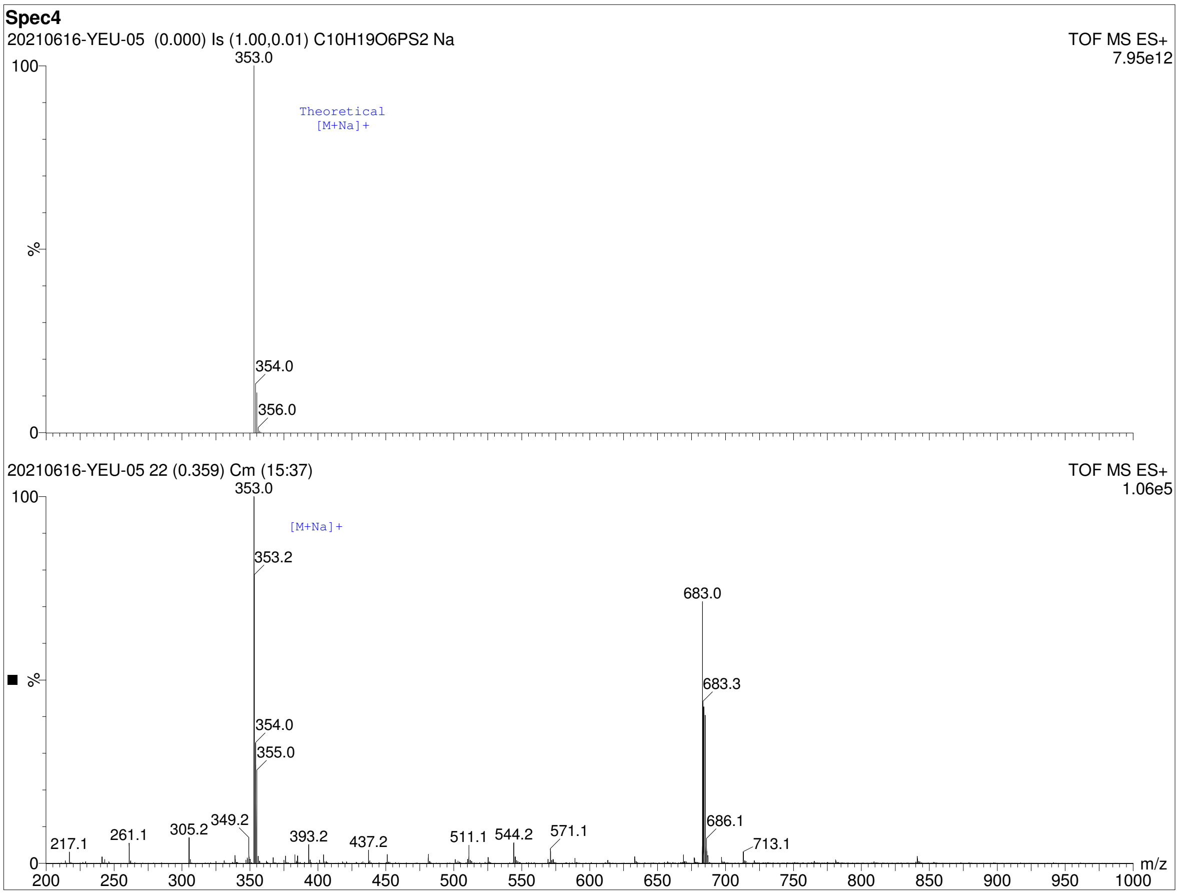


Figure 15: Mass spectrum of Spectracide (replicate 4) (Spectracide Catalog no. 071121309006) given by time-of-flight mass spectrometry. The theoretical mass spectrum of malathion is shown in the upper spectrum and the measured mass spectrum of Spectracide is shown in the lower spectrum.

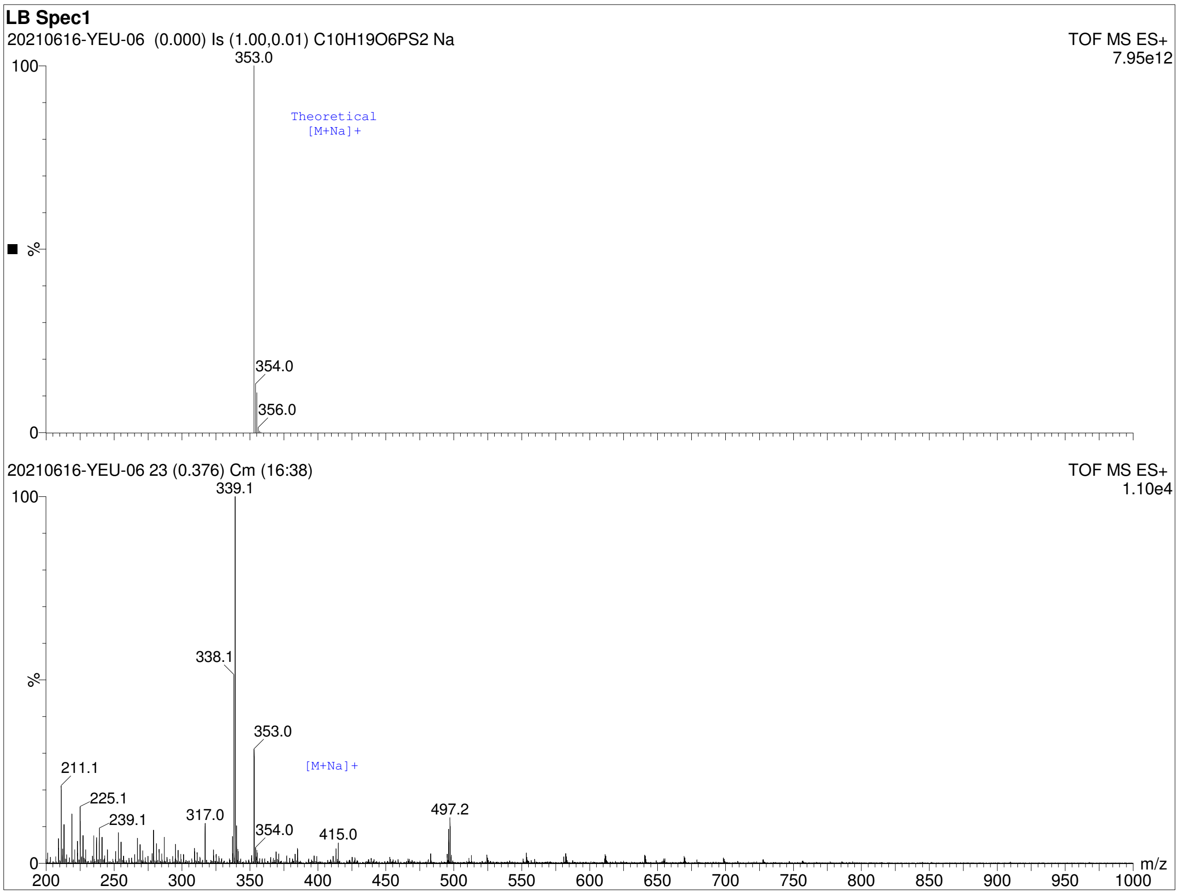


Figure 16: Mass spectrum of a 5% Spectracide in LB broth (replicate 1) given by time-of-flight mass spectrometry. The theoretical mass spectrum of malathion is shown in the upper spectrum and the measured mass spectrum of the solution is shown in the lower spectrum.

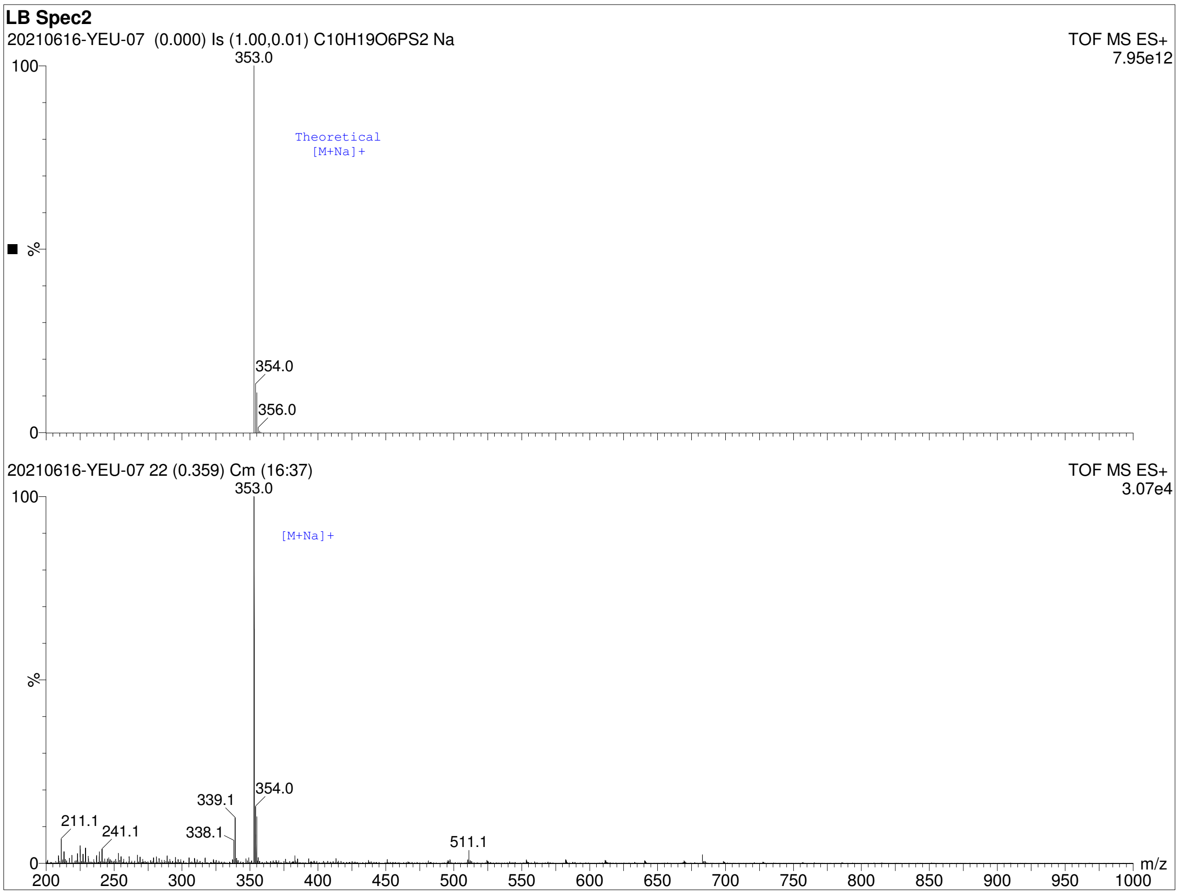


Figure 17: Mass spectrum of a 5% Spectracide in LB broth (replicate 2) given by time-of-flight mass spectrometry. The theoretical mass spectrum of malathion is shown in the upper spectrum and the measured mass spectrum of the solution is shown in the lower spectrum.

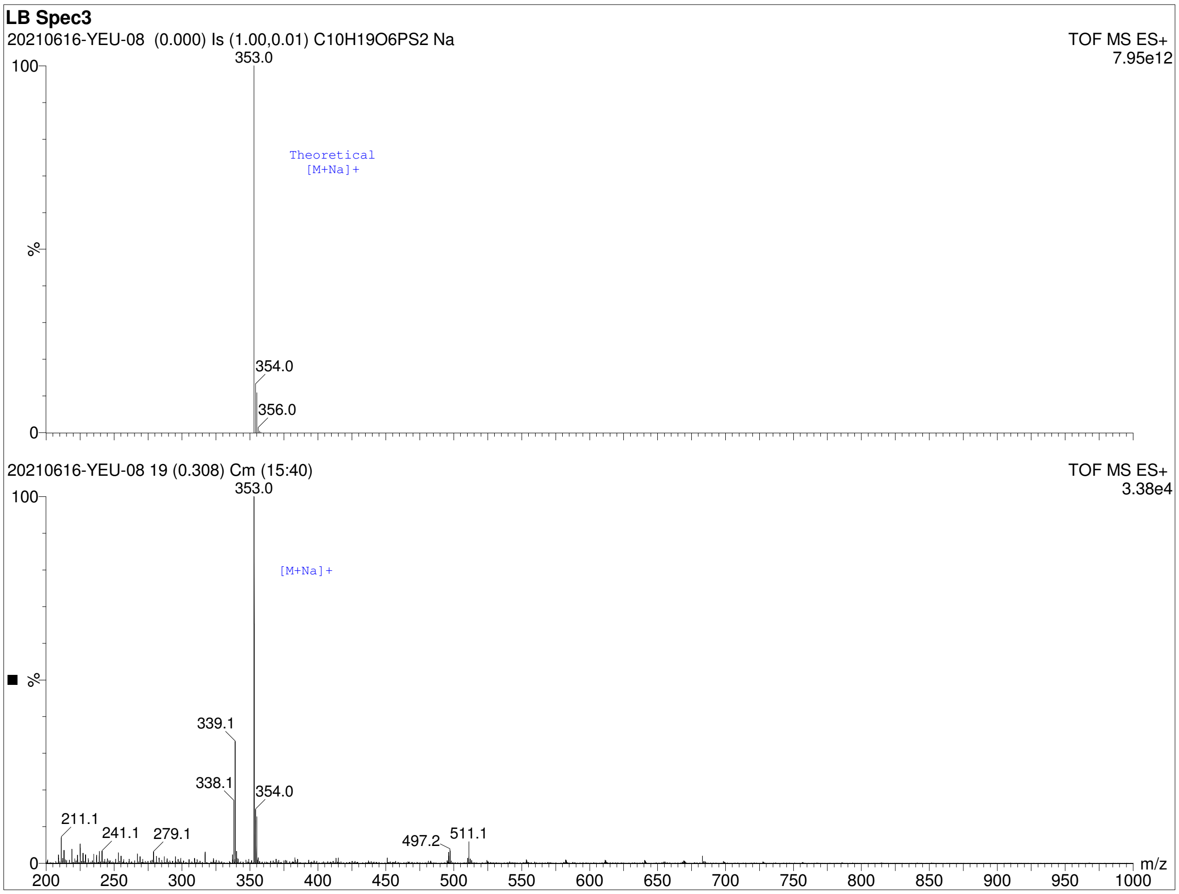


Figure 18: Mass spectrum of a 5% Spectracide in LB broth (replicate 3) given by time-of-flight mass spectrometry. The theoretical mass spectrum of malathion is shown in the upper spectrum and the measured mass spectrum of the solution is shown in the lower spectrum.

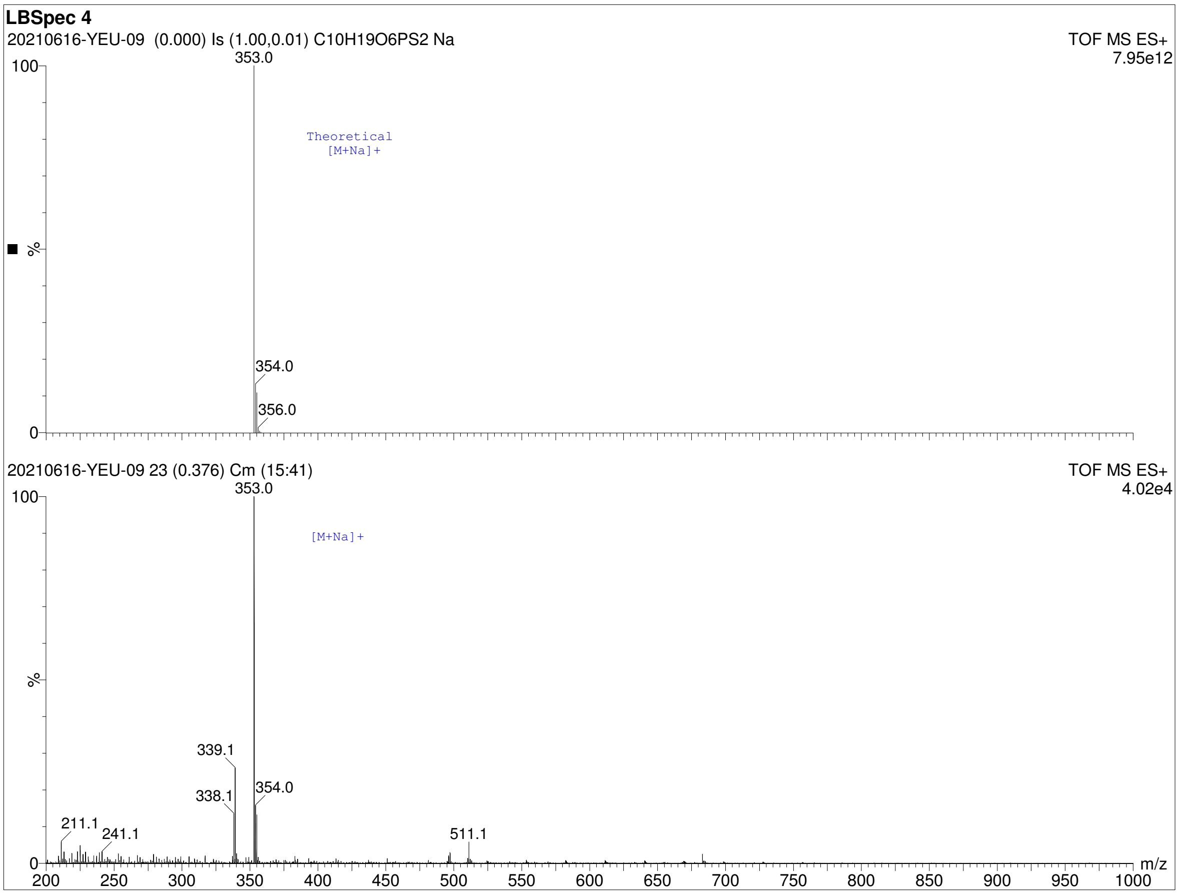


Figure 19: Mass spectrum of a 5% Spectracide in LB broth (replicate 4) given by time-of-flight mass spectrometry. The theoretical mass spectrum of malathion is shown in the upper spectrum and the measured mass spectrum of the solution is shown in the lower spectrum.

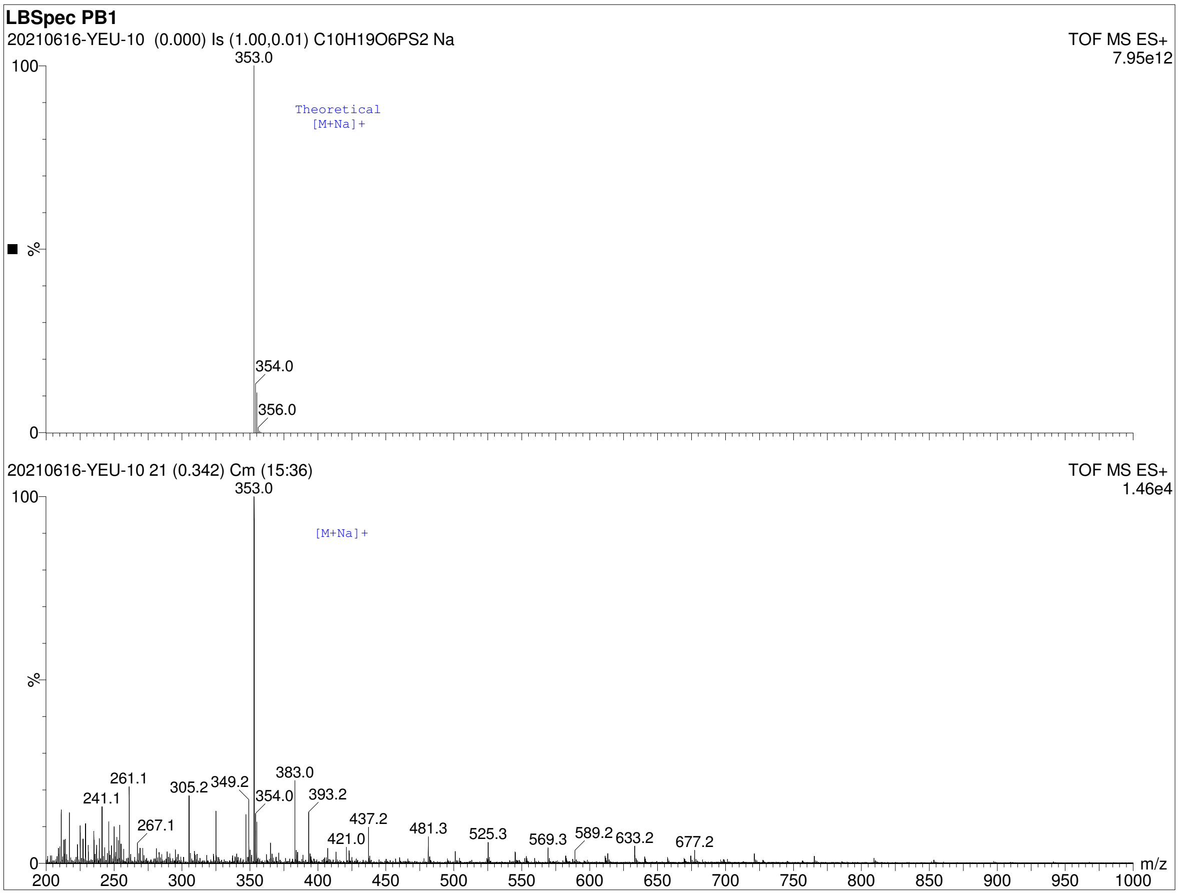


Figure 20: Mass spectrum of a 5% Spectracide in LB broth after photobleaching (replicate 1) given by time-of-flight mass spectrometry. The theoretical mass spectrum of malathion is shown in the upper spectrum and the measured mass spectrum of the photobleached solution is shown in the lower spectrum.

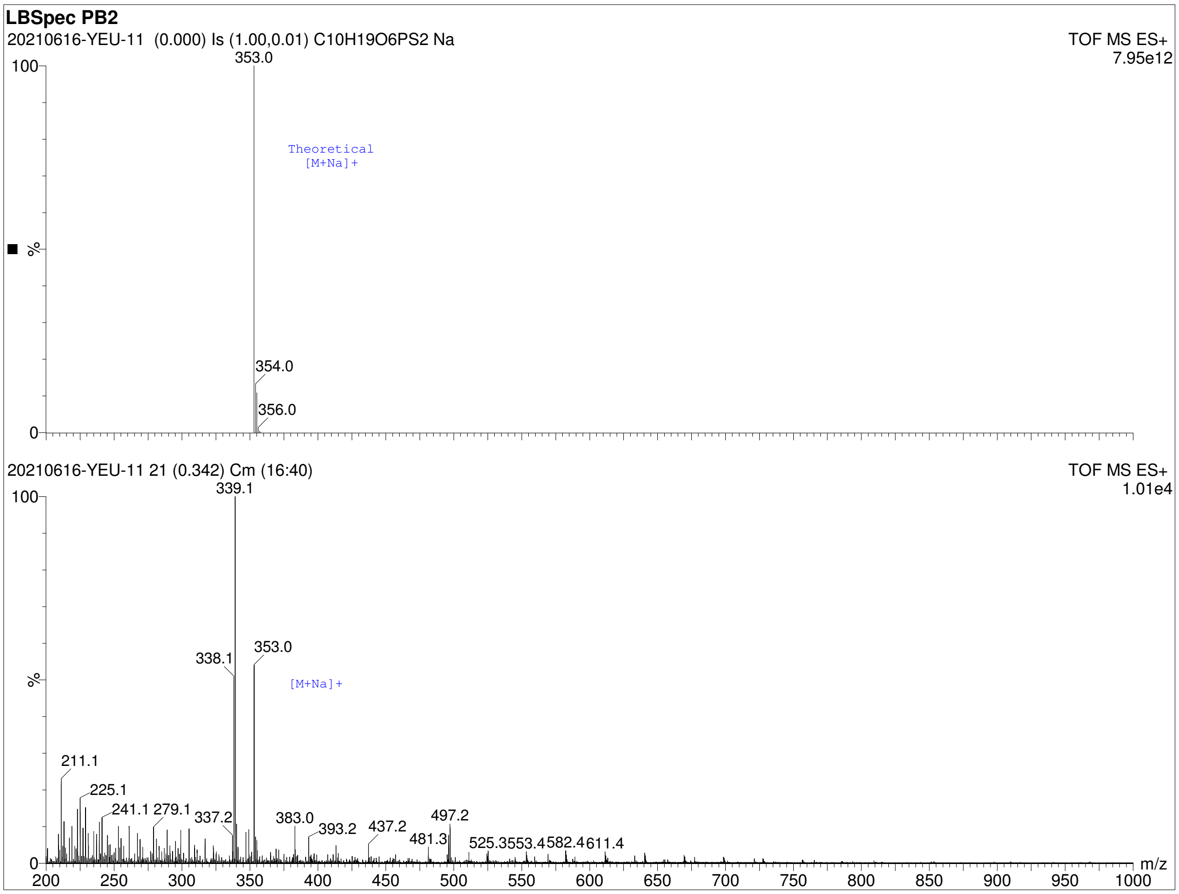


Figure 21: Mass spectrum of a 5% Spectracide in LB broth after photobleaching (replicate 2) given by time-of-flight mass spectrometry. The theoretical mass spectrum of malathion is shown in the upper spectrum and the measured mass spectrum of the photobleached solution is shown in the lower spectrum.

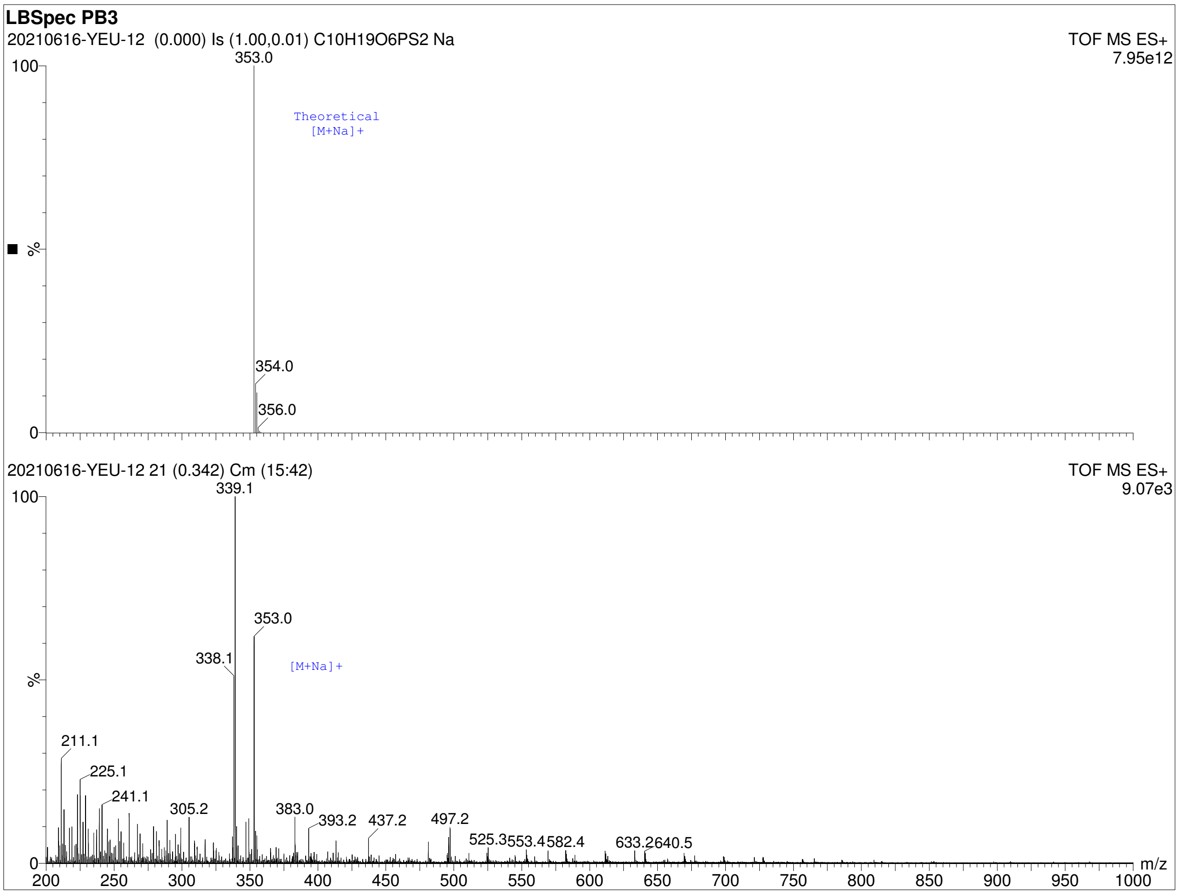


Figure 22: Mass spectrum of a 5% Spectracide in LB broth after photobleaching (replicate 3) given by time-of-flight mass spectrometry. The theoretical mass spectrum of malathion is shown in the upper spectrum and the measured mass spectrum of the photobleached solution is shown in the lower spectrum.

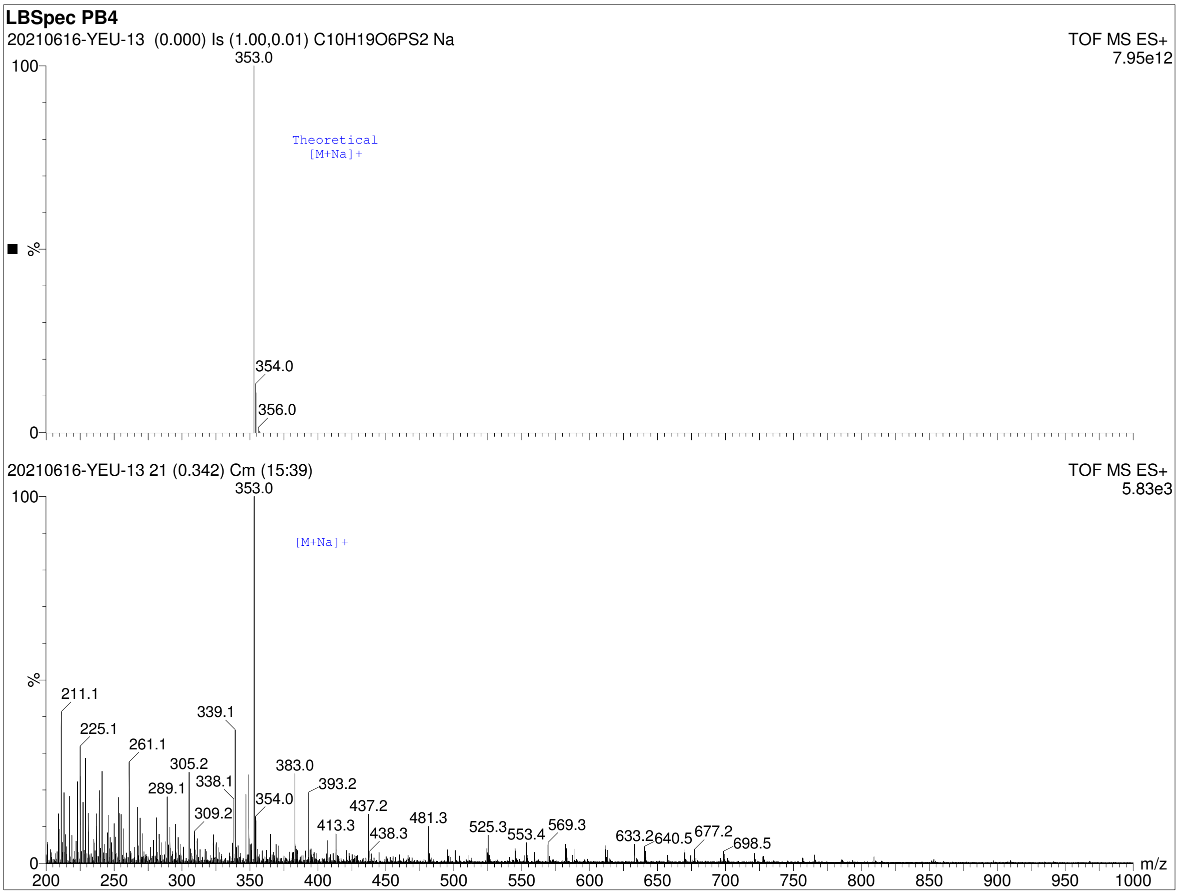


Figure 23: Mass spectrum of a 5% Spectracide in LB broth after photobleaching (replicate 4) given by time-of-flight mass spectrometry. The theoretical mass spectrum of malathion is shown in the upper spectrum and the measured mass spectrum of the photobleached solution is shown in the lower spectrum.

### 3 Supplementary Tables

| Time point (minutes) | $OD_{600}$ | Volume harvested (mL) | Malathion induction |
|---|---|---|---|
| 0 | 0.5 | 10 | |
| 10 | – | 10 | x |
| 20 | – | 10 | |
| 30 | – | 10 | |
| 40 | – | 10 | |
| 50 | – | 10 | |
| 60 | 1.0 | 10 | |
| 70 | – | 10 | |
| 80 | – | 10 | |
| 90 | – | 10 | |

Table 1: Metadata for the time-series RNAseq experiment.

| Sensor | Strand | Loci | Promoter sequence |
|---|---|---|---|
| PFLU_6124 | Antisense | 6709164 - 6709017 | AACATTTGCTTATGTAGCGCGTGATCGGAAATCACTAC<br>CCGGCAGTTGAATAGGGGCAGAACCGCCCCTATACTCT<br>GCGCGCATTTTGTCGGCACAAATTATGCCAAGTTATTG<br>ATTTCCGGCAGCCGACCATTGAGGAGCAAGAGTG |
| PFLU_0841 | Sense | 950865 - 951131 | TTCGCTTTACGTTCCACAAAAACGCCCAGCCTCCTCAC<br>GGAGCTGGGCGTTTTTTATTGCCTGCGATTTATACACA<br>AATTTCGCCGTGACAACTTGCCACATCCGTAGACCCCC<br>TATACTACAAGGCCTGGAGGCTGAGCCCAGGGCAATTC<br>CCTTGTCATACGTGGGGCTTTTCATTACCATTCGGCAA<br>AATTTTTATAAGTAAAGATTCAACACTTAGTAGACGCC<br>TGATTTAACAGGCCAAAAAAGCTGATGGGAGAGGACT<br>GA |
| PFLU_4736 | Sense | 5213048 - 5213188 | AGTGCTGGCAGAGGACGCTGGGTTTTTCTACACTGTGC<br>ACGAGATATTCCGTGCGCAGATTTATTGTCATTCGCGC<br>CTAAAGTTCGTCCGGGTATTGCCGAAAACATGGCAAGC<br>GTCCAAATACCCAGAGGTTTTTTGATC |
| PFLU_1823 | Sense | 1989934 - 1990137 | GCGAGATAATAAGAAACCACGGCGGAGTTGCCCGTCG<br>TGAGCCTTGCGCGCAAGACTCACCGCGGAATATCCGCT<br>GGACGCAGTCTTGCGCAGCTTTACGGGCCTTGAGCCCC<br>GCAAGCTGCGCAAGCAGCAGTCACAGGTGGCGCGGCA<br>CTCATAATGAGCGCAGCGCCGAATGCGCAGTACCTAAC<br>GAAGACGGTAAAAAGC |
| PFLU_3761 | Antisense | 4158693 - 4158135 | CTGTGACACGTCGCCAAGGCAGGCGCGGCGGATAGTT<br>TCAGTTCGGCGTCATACAAGTGCACTGCACCCCACTTC<br>ATCGTGGCCGTTTGCGAAAGCGATTGTCCGCTTGCGAC<br>GCGGCACAATCAGGGTATGTGCGCAGCTTGGCTTCCCA<br>GTAATTGCCCCATTAAATTTGTGGCTTTTCTGACGAGC<br>TTTTACTCGTCATCTCTTTGTTTTTTTACTATTATCGT<br>TCACCTGCGCACTCAAGGAAGAGAGGCTGAGCGCCTTG<br>AGGCTGGTAGAAAATTCATACTCGATCACTGAACGAGT<br>TATTGCTTTTACCCAGAACCTAACGACTCAGCCAACCA<br>TAAATACCTCTTGGTGAAACCGATGGATAAAATGTGTG<br>GCATCGTTGTAGTGGTAGGAC |
| PFLU_5502 | Antisense | 6038217-6038089 | GTGTGATCCGCTTGAAGCCCGGCAGCTAGTGCGCTGCC<br>GGGTTGATTATTTGTTATTACAGCGATATTATCTCGCG<br>CCCTATTTCTTGGCTTCCGGGGCGTAGGTAGCTGTCAA<br>TTGGAGTCCCACTGA |

| | | | |
|---|---|---|---|
| PFLU_1836 | Antisense | 2003829 - 2003581 | CGCCGCGCCATCAGCCAACTCCGACTGGCGTGAAAGAC<br>GAAAGTGCGGCAGTCTTAGGCACCCGAACGGGCCCAT<br>AAACAGGCCCGGTTTAAAATTTCAGTGAACAAGTGTA<br>CATTCAGTACCTTGCCGCTGTGACTTTCACTACAACGC<br>AATAGTCTATGTGTAGGCTGCCGACATGAGGCATGAAC<br>GCTTCATTCGGTTCGGGAAGATTTGCCCTACCCTGCCG<br>CATGGGATTATTGAGGAGCTCGC |
| PFLU_0376 | Sense | 417961 - 418174 | AAGTCATAACTGCTTACACATCAACCGGTTGCCGGTAC<br>TCCTCTGCGTAAGTGTCTGCCCCTGAGCTTTGCCGCAC<br>CGATGTGGGGCTTTTCCGACATATGCCGATAACAAATA<br>GCCGTGAAACCTTTGTGACGAGCAACGAGTGGGTTAG<br>GATCGCACCCCGAAATGGTGCAGCCCTTTTGCGCGCCG<br>ACCTTACAAAAATCGTTCAGGGGAC |
| PFLU_1815 | Antisense | 1980804 - 1980440 | GGCTTTTTTTCACACTGAAGAGCCCCTAACAATCAGGG<br>CAAAGTTGTTGGGGAGTGCGACTGGTCAGGTAAGCAC<br>CACCCAGGGAGTGCGACCCCCAATGAAAGCAAGCCCA<br>AAAGCCCTTGGCGGTCGGTGGCCGAGTATAGACAGTT<br>AGGTTACTAATGACAACGCGCACTCCTCACCCTAATAG<br>CTGATTGCGCTGGCGGGATAAAAGGCGTAAATGGCGC<br>TCAATTTCGAGGAAAAAGTACGGTTAAAGCCTTCTGGG<br>GCAAGACTTTAGGCAAATTGACATCTGAATTTATCTCA<br>CTATAGTGGTGCGGGCCCTGCGTGGGGGGTCTGTCTG<br>ATGATTTGAAGCATAAATAGGAGGCCAC |
| PFLU_0953 | Sense | 1058342 - 1058453 | TGGAATGTATCAGGGCTATGAAGGTGATTGGTGTTCA<br>GCAAAGGTCTGGTCTGCTATTATCGCCAGCCTTTGTTG<br>ATACCAGTTCGCAATTTGCGCTGAAGCGGTCCAAGCC |
| PFLU_1380 | Antisense | 1527252 - 1526967 | GGCAGTAAAACCTCAATCAGGACACTGGGGGCTATCG<br>TTAACGCAACGTTAATAGACGTAAACGATCATCCGAAT<br>ATTTGTGGGACGACACCGTCATGGGTGCCGAACGTAAT<br>GGAATCGAGGCTTCGGGCGTTGCTTTGTCAACACTCCG<br>CGAAGCCTGTCAAGAGTTTACAAACAACCATGAACGTA<br>AGTATATTGCGTAGCAAGCTACTTATCCACTCACAGCT<br>TGTTTTTTACCCTTCCACACTTCTTGTGCGCACCCTGC<br>GCGCCTGACCCGAGGATCTTC |
| PFLU_4612 | Antisense | 5088655 - 5088549 | TGCGTGGCAAATATCTCTTACGTGTAGGCAAGTTCTGT<br>TAGACTTGTCGCCGAGTTGTCCCCCGGTTTGTGGGACT<br>GCTTTACAATCACCAGATGGGGATTTAACGG |
| PFLU_4150 | Sense | 4592631 - 4592843 | GATTTGCCGCTGATCTCACGGCTTTTTTGGCGGTAAAA<br>CAGGCTTAAAACTGCCGCTTCTCACAAATTACGCAGCT<br>TTTACGGCTTTTTTTACCAGTTGATATTTCGAGCCAAA<br>GCCCCGTAAATCGGGGCTTTCAGCCGGATCAGGCACTA<br>AACGCAACGCTATTGATTAGCAAACAATGCCTTGGGGG<br>GGCTCCCAAGCCGAACATTTCGACTATGATAGCCCGGT<br>GTGCCCAGTTGGCCTGAGCAGCACAGCACTACTGAAAA<br>TATATGTTTCTTGGAGATACACC |
| PFLU_1302A | Antisense | 1440968 - 1440759 | GGCGGGTGTCCTGAATAGCGAGGTGGAAAAATACTGT<br>GGGGCATCTTACCGGGGCCGGCGTTGGGGTTCAAAGG<br>TCACAGGGCTTTTCTTGATGAATGCGCCGGCGGCTATA<br>AGCCGCAGCCAGCGGGGCCGGGGTTAATATTGCCGCG<br>CAGGGCGACCGTGGATACCACCGTCAGTCACGAATTTA<br>GAGAAACCTTCGGAAATACCATTGGCACGTTCCGGAAA<br>AAGGGTTAAGGTGGCGCCACTGTGCTGCTTGTGTCACT<br>GAGAATCTCTACACGATATGTTGAATTTCGATCCAACC<br>ATCTCCAAGAATTTTTCCTGCTCTTTGCACTCAGTCTC<br>GGCCAGGGCTTTTCCTGAGTCGCAGTTAACTTTGTCCA<br>AGGAGATACACC |

| | | | |
|---|---|---|---|
| PFLU_1358 | Sense | 1498195 - 1498311 | AACAGCCTGCATCCATTGATGCAGGTCAGTTATTGCCC<br>TTCTTTACGCTCCGTCGTGGGCGACATTGATCCCCGTC<br>AATTTTCCAATCCGCCTTCTGCATTAACTTAGCCCTAT<br>CGCAACAGGGCAAGTGCAGGAGGCCGGTC |

| Part | Sequence |
|---|---|
| BCD1 | GGGCCCAAGTTCACTTAAAAAGGAGATCAACAATGAAAGCAATTTTCGTACTGAAACATCT<br>TAATCATGCACAGGAGACTTTCT |
| ECK120033736 | AACGCATGAGAAAGCCCCCGGAAGATCACCTTCCGGGGGCTTTTTTATTGCGC |
| sfGFP | ATGCGTAAAGGCGAAGAGCTGTTCACTGGTGTCGTCCCTATTCTGGTGGAACTGGATGGT<br>GATGTCAACGGTCATAAGTTTTCCGTGCGTGGCGAGGGTGAAGGTGACGCAACTAATGGT<br>AAACTGACGCTGAAGTTCATCTGTACTACTGGTAAACTGCCGGTACCTTGGCCGACTCTGG<br>TAACGACGCTGACTTATGGTGTTCAGTGCTTTGCTCGTTATCCGGACCATATGAAGCAGCA<br>TGACTTCTTCAAGTCCGCCATGCCGGAAGGCTATGTGCAGGAACGCACGATTTCCTTTAAG<br>GATGACGGCACGTACAAAACGCGTGCGGAAGTGAAATTTGAAGGCGATACCCTGGTAAAC<br>CGCATTGAGCTGAAAGGCATTGACTTTAAAGAAGACGGCAATATCCTGGGCCATAAGCTG<br>GAATACAATTTTAACAGCCACAATGTTTACATCACCGCCGATAAACAAAAAAATGGCATTA<br>AAGCGAATTTTAAAATTCGCCACAACGTGGAGGATGGCAGCGTGCAGCTGGCTGATCACT<br>ACCAGCAAAACACTCCAATCGGTGATGGTCCTGTTCTGCTGCCAGACAATCACTATCTGAG<br>CACGCAAAGCGTTCTGTCTAAAGATCCGAACGAGAAACGCGATCATATGGTTCTGCTGGA<br>GTTCGTAACCGCAGCGGGCATCACGCATGGTATGGATGAACTGTACAAATGATGA |
| 5' overhang | GAACGGTCTCAGCAT |
| 3' overhang | GTCGTGAGACCTTACG |

Table 3: Sequences for the parts used in the reporter cassette.

| Malathion reporter | Locus tag | Time point (hours) |
|---|---|---|
| atpB | PFLU_6124 | 1.0 |
| petA | PFLU_0841 | 2.0 |
| anti-sigma 28 factor | PFLU_4736 | 3.2 |
| sucC | PFLU_1823 | 8.1 |
| Uncharacterized protein I | PFLU_3761 | 12.9 |
| rpoA | PFLU_5502 | 15.0 |
| fabA | PFLU_1836 | 14.0 |
| Putative ABC transport protein | PFLU_0376 | 0.9 |
| gltA | PFLU_1815 | 3.2 |
| lpxC | PFLU_0953 | 0.7 |
| acrA | PFLU_1380 | 3.1 |
| Putative outer membrane porin A protein | PFLU_4612 | 2.0 |
| cspA2 | PFLU_4150 | 2.4 |
| capB | PFLU_1302A | 8.5 |
| Uncharacterized protein II | PFLU_1358 | 5.6 |

Table 4: The time points at which the Hill functions are fit to each reporters' response.
